# A novel molecular interaction in bronchiolitis obliterans syndrome in lung transplantation patients: the role of SERPINA3 and osteoprotegerin

**DOI:** 10.1101/2025.10.27.684757

**Authors:** Yanzhe Liu, Eline A. van der Ploeg, Theo Borghuis, R. Ian Menz, Wim Timens, Judith M Vonk, Barbro N Melgert, C. Tji Gan, Janette K Burgess

**Affiliations:** University of Groningen, University Medical Centre Groningen, Department of Pathology and Medical Biology, Groningen, The Netherlands; University of Groningen, University Medical Centre Groningen, Groningen Research Institute for Asthma and COPD, Groningen, The Netherlands; University of Groningen, University Medical Centre Groningen, Department of Pulmonary Medicine, Groningen, The Netherlands; School of Life Sciences, Faculty of Science, University of Technology Sydney, Sydney, NSW, Australia; University of Groningen, University Medical Centre Groningen, Department of Epidemiology, Groningen, The Netherlands; University of Groningen, Department of Molecular Pharmacology, Groningen Research Institute for Pharmacy, Groningen, The Netherlands

**Keywords:** Lung transplantation, chronic lung allograft dysfunction, bronchiolitis obliterans syndrome, TNFRSF11b, fibrosis, airway obstruction, airway remodeling

## Abstract

**Background and objective:** Chronic lung allograft dysfunction, significantly limits survival after lung transplantation. The obstructive phenotype bronchiolitis obliterans syndrome (BOS) is characterized by the abnormal activation of epithelium in airways accompanied by fibrotic changes with excessive deposition of extracellular matrix, resulting in narrowing and ultimately obliteration of airways. Mast cells, characterized by mediators such as tryptase and chymase, play a role in lung fibrosis. The proteins, osteoprotegerin (OPG) and SERPIN family member A 3 (SERPINA3) have previously been associated with progression of lung fibrosis. Tryptase-mast cells can produce OPG, while chymase released from mast cells interacts with SERPINA3. This study aimed to investigate if and how SERPINA3, OPG and tryptase/chymase-positive mast cells are related to fibrotic airway obliteration, and potentially show an association with BOS severity.

**Methods:** SERPINA3 levels in serum from patients with or without BOS were examined using ELISA. In lung tissue from patients with BOS, we identified non-cartilaginous airways and classified them into normal, partially and completely obstructed airways. Immunohistochemistry was used for detection of SERPINA3, OPG, chymase and tryptase. Colocalization of, and interactions between, SERPINA3, OPG and tryptase were assessed by immunofluorescence, proximity ligation assay, and AlphaFold modeling.

**Results:** SERPINA3 levels in serum from BOS patients were higher compared to non-BOS patients. A low percentage of SERPINA3 and OPG was detected in partially and completely obstructed airways. Cells positive for OPG and SERPINA3 colocalized with tryptase-mast cells in airways from BOS lung tissue. OPG colocalized with SERPINA3 and their staining positively correlated in partially obstructed airways. In completely obstructed airways, OPG, SERPINA3 and tryptase staining areas all positively correlated with each other.

**Conclusion:** SERPINA3 and OPG are expressed by mast cells and these proteins may form a complex in lung tissue with potential for contributing to airway remodeling in BOS.

## INTRODUCTION

Chronic lung allograft dysfunction (CLAD) limits survival after lung transplantation^1^. Bronchiolitis obliterans syndrome (BOS) is the most common form of CLAD, affecting 50% of patients five years after lung transplantation^2^. BOS is a clinical diagnosis based on persistent lung function decline of at least 20% of forced expiratory volume in one second (FEV1) compared to baseline after lung transplantation^3^. Bronchiolitis obliterans is defined as a histopathological hallmark of BOS. It is characterized by an abnormal repair process in predominantly non-cartilaginous, smaller airways, which includes initial epithelial inflammation with subsequent fibrosis formation. Excessive deposition of extracellular matrix^4^ in the submucosa of airways then ultimately contributes to complete obstruction with the absence of any remaining epithelium in smaller airways^5^. Such fibrotic abnormalities result in partial or complete obliteration of the airway lumen, leading to a decline of lung function and symptoms of breathlessness^6^. However, the mechanisms underlying bronchiolitis obliterans remain poorly understood.

Mast cells play a critical role in abnormal repair processes by releasing mediators that drive extracellular matrix remodelling^7^. Tryptase which is released from mast cells, is also recognized as a marker to identify mast cells^8^. Chymase, another mediator secreted from a subset of mast cells, plays a crucial role in orchestrating extracellular matrix degradation by regulating the activation of matrix metalloproteases to maintain tissue homeostasis^9^. Chymase interacts with SERPIN family member A3 (SERPINA3), also known as alpha-1-antichymotrypsin, to form an irreversible chymase-SERPINA3 complex which inhibits chymase activity^10^. We recently reported that SERPINA3 was higher in serum from patients with BOS after lung transplantation compared to non-BOS patients^11^. SERPINA3 is an acute phase protein that is secreted by the liver in response to inflammation^12^ and has been linked to fibrotic responses in various tissues. In a model of bleomycin-induced lung fibrosis, high levels of SERPINA3 were found and silencing an isoform of SERPINA3 significantly reduced expression of collagen type I and fibronectin^13^. Furthermore, *SERPINA3* gene expression positively correlated with expression of collagen type II A1 mRNA in cartilage and its silencing downregulated expression of genes related to assembly of collagen fibrils in mesenchymal stem cells^14^. Whether disruption of the relationship between SERPINA3 and chymase is an important driver of the fibrotic extracellular matrix changes observed in BOS lung disease has yet to be examined.

Osteoprotegerin (OPG), a member of the tumor necrosis factor (TNF) receptor superfamily, is a glycoprotein that plays a key role in regulating homeostasis of bone extracellular matrix^15^. Beyond its traditional function in bone, OPG has increasingly been implicated in various fibrotic conditions. Elevated serum OPG levels, along with its significant presence in lung tissues in patients with idiopathic pulmonary fibrosis (IPF) have been observed. Elevated serum OPG was also associated with decreased lung function and progression of IPF^16^. Moreover, a recent study has identified OPG as a marker of early fibrosis and an indicator of responsiveness to anti-fibrotic therapy^17^. Importantly, elevated levels of serum OPG were strongly associated with graft failure and related to all-cause mortality in renal transplantation recipients^18,19^. Of interest, tryptase-positive mast cells are capable of secreting OPG^20^ to regulate bone turnover.

Building on these findings, this study aimed to investigate if SERPINA3, OPG and chymase/tryptase-positive mast cells were related to airway obliteration, potentially contributing to BOS severity. We hypothesized that enhanced OPG production by tryptase-positive mast cells interacts with the increased SERPINA3 to contribute to the airway obstruction of BOS. To investigate this hypothesis, we identified multiple non-cartilaginous airways with varying degrees of submucosal fibrosis in lung tissue from patients with BOS after lung transplantation to detect the presence of SERPINA3, OPG and chymase/tryptase-positive mast cells in relation to fibrosis progression in BOS.

## MATERIAL AND METHODS

### Ethics information

For serum analysis, lung transplant (LTx) patients who underwent bilateral lung transplantation between 2004 and 2017 in the University Medical Center Groningen were screened. BOS patients who progressed to stage three according to the International Society of Heart and Lung Transplantation Guidelines were selected if longitudinal serum samples were available. BOS patients were matched to non-BOS patients for sex, age at LTx, diagnosis necessitating LTx, immunosuppression and storage time of the samples (Table S1). Explants from LTx patients who received a re-transplantation due to BOS or deceased with end stage BOS were collected for tissue analysis. All patients received immunosuppression according to protocol (Table S2). Patients provided written informed consent for use of material. The study was approved by the medical ethics committee of the University Medical Center Groningen (METc 2014/077, METc 2021/610, research register number: 202000737), adheres to the UMCG Biobank Regulation and was conducted in accordance with the WMA Declaration of Helsinki and Declaration of Istanbul. Patients were enrolled in the ongoing, prospective TransplantLines Biobank and Cohort Study (ClinicalTrials.gov identifier: NCT03272841), in which, since June 2015, all (potential) solid organ transplantation patients and (potential) living organ donors (aged ≥18 years) at the University Medical Center Groningen (UMCG, The Netherlands) have been invited to participate.

### Patient information

The patient cohort used in this study for the detection of SERPINA3 in serum was the same as that used in our previous proteomic study^21^. Briefly, non-BOS patients (n=19) were matched to BOS patients (n=19). Serum samples were identified that had been collected at 12, 6, 3 months before BOS onset, and subsequently at BOS stage 2 and BOS stage 3. We also matched the time points of collection of samples for the non-BOS patients to BOS patients. The information relating to this cohort for the serum SERPINA3 analysis is in Supporting information Table S1, and the cohort used for detection of SERPINA3, tryptase, OPG and chymase in lung tissue is in Supporting information Table S2.

### Soluble SERPINA3 detection

ELISA was used to assess the level of SERPINA3 in serum according to the manufacturer’s instructions, Human alpha 1-Antichymotrypsin ELISA Kit (ab157706, Abcam, Cambridge, United Kingdom) (details in supporting information)

#### Lung tissue staining

Structures within lung tissue were identified using hematoxylin and eosin (H&E), Verhoeff’s, and Martius Scarlet Blue (MSB). SERPINA3, tryptase, chymase and OPG were detected in BOS lung tissue using immunohistochemistry. To investigate (co)localization of above proteins in BOS lung tissue we used immunofluorescence. Protein-protein interactions were studied using proximity ligation assay (NB.MR.HRP.100, Navinci, Uppsala, Sweden). Full details are provided in Appendix S1-ADDITIONAL METHODS (Supporting information).

### Protein interaction modelling

To predict biomolecular interactions between SERPINA3, tryptase, chymase, and OPG we used AlphaFold Server^22^ (https://alphafoldserver.com). The details are described in Appendix S1-ADDITIONAL METHODS (Supporting information).

### Airway categorization, identification and annotation

All airways in H&E-stained lung tissue sections from patients with BOS (n=6, 3-5 sections per patient) were identified by an experienced lung pathologist (WT) and all airways without presence of cartilage and submucosal glands were included. For each airway, the airway region of interest was defined as the region between the outer border of the adventitia of the airway to the inner border of the airway lumen for normal and partially affected airways. When airways were completely obstructed, the circumference of the outer border of the adventitia was identified as the boundary of the region of interest. All airways were also identified in SERPINA3, tryptase, chymase and OPG-stained sections (3-5 sections per patient / per protein of interest) and categorized according to the classification code established in the H&E-stained sections. All details are summarized in Appendix S1-ADDITIONAL METHODS (Supporting information).

### Image analyses

All stained slides were scanned with a Nanozoomer (Hamamatsu Photonic K.K., Japan). NanoZoomer Digital Pathology Image (NDPI) files were first converted to TIF images in Aperio ImageScope (Leica) and subsequently processed in Adobe Photoshop 2024 (Adobe Inc. California, United States) to exclude artifact areas such as folded tissues and carbon-based pigments. Next, Image J win 64 software ^23^ was used to quantify areas with positive staining. All images were split into blue (hematoxylin-image), and red (Nova Red-image) pixels using the color deconvolution plugin of Image J^24^. To calculate the total amount of tissue, images were converted to 8-bit grey scale. The total number of pixels that represented the total area occupied by tissue versus the number of pixels that were positively stained for each protein of interest within the tissue area of interest were identified using the threshold feature of Image J. Percentage of positive area within the whole tissue region was then calculated. Data analyses were performed with R software 4.4.0 (Boston, Massachusetts, USA). All details are provided in Appendix S1-ADDITIONAL METHODS (Supporting information).

### Statistical Analyses

Patient characteristics are described as mean ± standard deviation for normally distributed data, or median [interquartile range] for non-normally distributed data. Data were tested for normality with a Shapiro Wilk test and visually assessed by histograms and P-P plots. If not normally distributed, data were Ln transformed to attain a normal distribution. If this did not yield a normal distribution, nonparametric tests were used when comparing two groups, i.e. Mann-Whitney U or student’s t-tests depending on normality of the data. Longitudinal data for serum SERPINA3 levels were examined with a linear mixed model using IBM SPSS Statistics 28.0.1.0 after Ln transformation of SERPINA3 levels for normalization, with intercept per patient as random effect. Differences in the percentages of positively stained area within sections of lung tissue between normal airways, partially and completely obstructed airways were assessed using linear mixed effects regression analysis in SPSS with two random effects on intercept, to correct for both per subject and paraffin block variability. For staining analysis natural log (ln) transformation was performed to achieve normality of the data. Correlations between SERPINA3, OPG and tryptase were assessed using linear regression with either Pearson or Spearman test depending on normality of the data. Correlation data are shown with 95% confidence bands of the best-fit line. p <0.05 was considered significant for all tests.

## RESULTS

### Higher serum SERPINA3 level in patients with BOS

SERPINA3 levels in serum were significantly higher in patients with BOS compared to non-BOS patients at BOS stage 1 and BOS stage 3 (Figure 1). SERPINA3 levels in serum correlated negatively to FEV1 in liters (estimate –0.421, 95% CI –0.695 - −0.135, p<0.005). In the twelve months before onset of BOS (12 months before BOS until BOS stage 1) LnSERPINA3 levels decreased in non-BOS patients per day before BOS onset, while LnSERPINA3 increased in BOS patients per day before BOS onset (−0.015 (95%CI –0.29 - −0.001) vs 0.01 (95%CI 0.005 - 0.044), p=0.01) (Figure S3).

**Figure 1.**
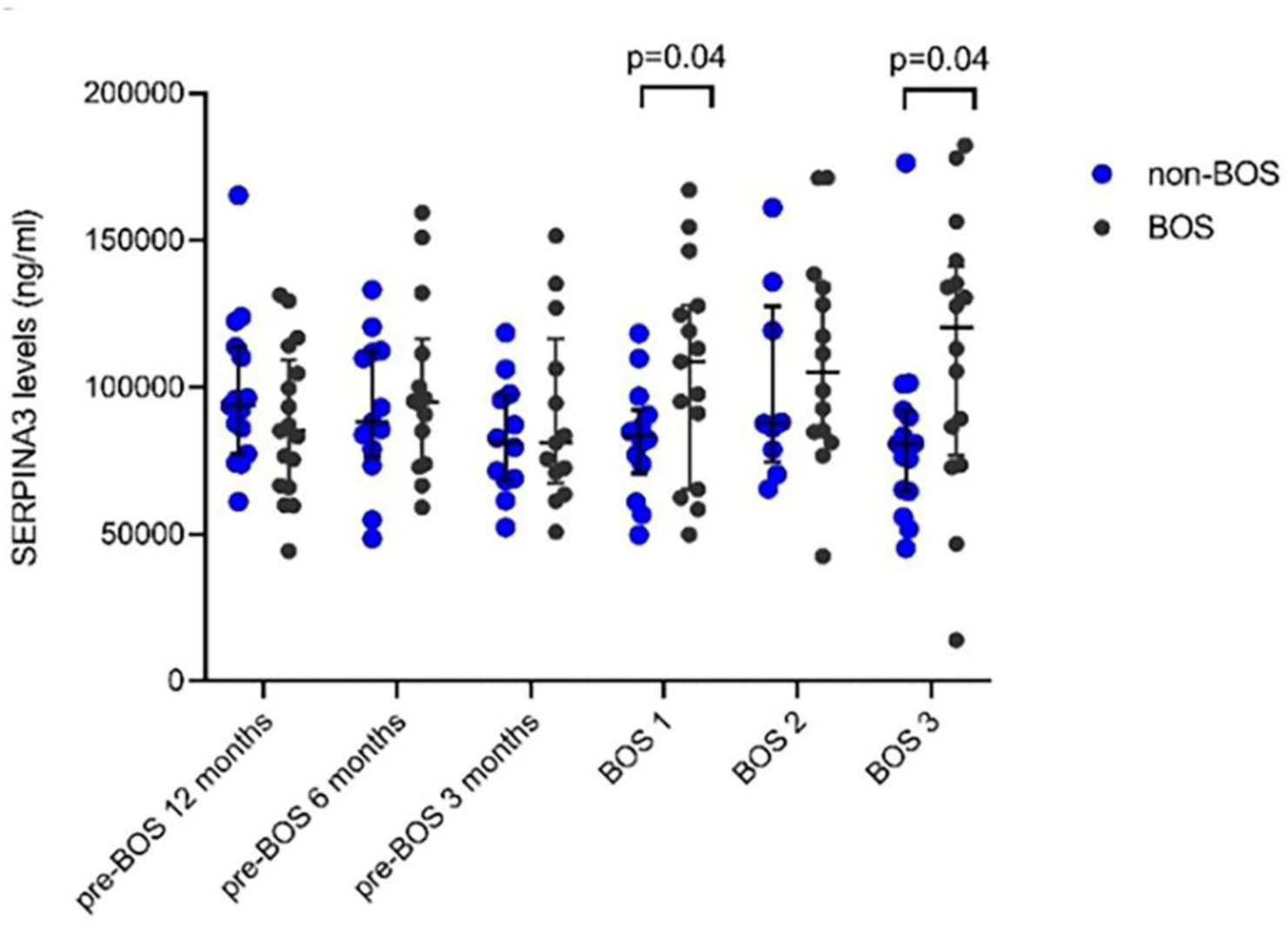
Serum SERPINA3 levels in patients with and without BOS. Serum samples from patients who did (n=19) or did not (n=19) develop BOS, from different time points relative to BOS diagnosis in the affected patients, as analyzed by ELISA. All points represent, at each timepoint, measurements from individual lung transplantation patients. Error bars reflect median [interquartile range]. Groups were compared at each time point using a Mann-Whitney U, p<0.05 was considered significant. Abbreviations: BOS; bronchiolitis obliterans syndrome.

### Low SERPINA3 deposition in partially and completely obstructed airways colocalizes with mast cells in the lung tissue from patients with BOS

The elevated serum SERPINA3 levels in patients with BOS prompted us to further investigate the localization of SERPINA3 in lung tissues from patients with BOS. Therefore, we used immunohistochemistry to identify SERPINA3. As it was not possible to access lung tissue samples from lung transplant patients without BOS to use as controls, we used SERPINA3 distribution in unaffected/normal airways in lung tissue from patients with BOS to compare to partially and completely obstructed airways in the same tissues (Figure 2A). We observed that in normal airways SERPINA3 localized in the adventitia and was specifically detected in a population of cells with a distinctive round morphology. In partially obstructed airways, the cells positively stained for SERPINA3 were mainly observed around the smooth muscle area and partly in the obstructed connective tissue area of the airway. When airways were completely obstructed the SERPINA3 staining was more prominantly present in the connective tissue in the luminal obstruction (Figure 2A). Subsequently, we quantified the percentage area of tissue with positive SERPINA3 staining within the whole airway wall excluding epithelium. The ln-transformed data indicated a lower percentage area positive for SERPINA3 in both partially and completely obstructed airways than in normal airways, which is likely mainly the result of the increase in connective tissue present in the scarred obstructed area. (Figure 2B).

**Figure 2.**
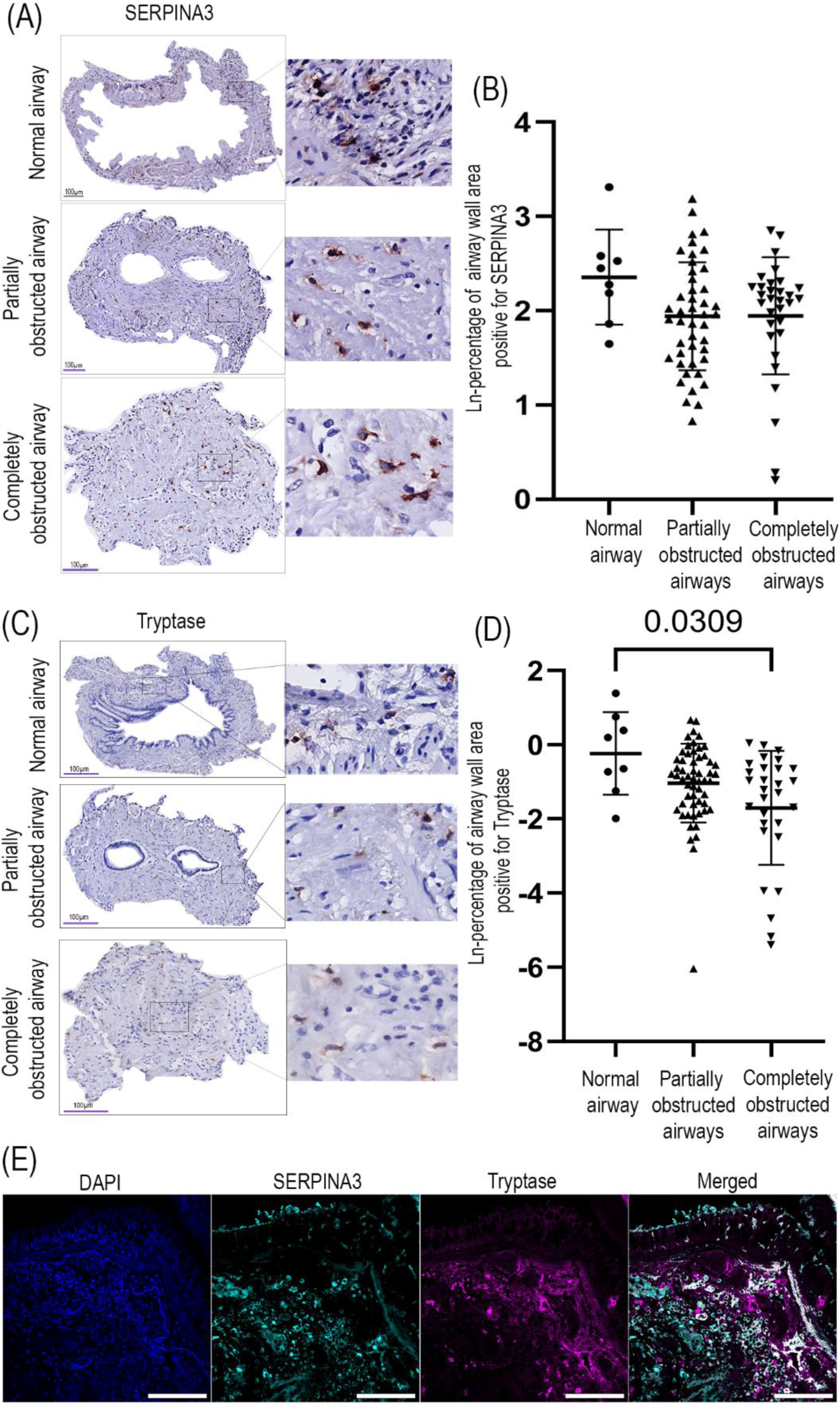
Localization of SERPINA3 and tryptase-positive mast cells in airways in BOS lung tissue. Formalin-fixed paraffin-embedded lung tissue from patients with BOS stained for SERPINA3 using immunohistochemistry. (A) SERPINA3-positive signal is shown in red as detected by Nova Red; nuclei are shown in blue as detected with hematoxylin. Sections were imaged at original objective magnification 400X, and digital magnified images are shown next to each representative image. (B) The percentage of total tissue area positive for SERPINA3 in normal, partially and completely obstructed airways is presented as ln-transformed values and each datapoint represents an individual airway. Data are presented as mean +/- SD and groups were compared using a linear mixed effects regression analysis (n= 6 patients, number of airways per patient: 0-6 for normal airways, 0-21 for partially obstructed airways, 0-23 for completely obstructed airways). (C) Formalin-fixed paraffin-embedded lung tissue from patients with BOS stained for tryptase using immunohistochemistry. Tryptase-positive mast cells localized to normal airways, partially and completely obstructed airways. (D) The percentage of total tissue area positive for tryptase in normal, partially and completely obstructed airways and is presented as ln-transformed values with each datapoint representing an individual airway (n= 6 patients, number of airways per patient: 0-6 for normal airways, 0-21 for partially obstructed airways, 0-19 for completely obstructed airways). Data are presented as mean +/- SD and groups were compared using linear mixed effects regression analysis, p<0.05 was considered statistically significant. (E) Colocalization of SERPINA3 and tryptase in lung tissue from patients with BOS stained for SERPINA3 and tryptase using immunofluorescence. Cell nuclei, SERPINA3, and tryptase are respectively shown in blue, cyan, and purple, colocalized regions are represented in white in the merged image (scale bar = 100 µm).

The cells positively stained for SERPINA3 were number-and distribution-wise suggested to be mast cells. To confirm this supposition, we stained for tryptase, which is only expressed by mast cells. In lung tissue sections from patients with BOS, we observed tryptase expression in all categories of airways with similar distribution as SERPINA3 (Figure 2C). Next, we calculated the percentage of staining area for tryptase-positive mast cells within each entire airway. We found that tryptase-positive mast cell area was significantly lower in obstructed airways compared to normal airways (Figure 2D). To explore whether SERPINA3-stained cells were mast cells, colocalization between SERPINA3 and tryptase-positive mast cells was determined by immunofluorescence in the airways of BOS lung tissue. We observed SERPINA3 and tryptase colocalized, indicating the SERPINA3-positive cells were probably tryptase-positive mast cells in the small airways from transplant patients with BOS (Figure 2E).

### More chymase-positive mast cells in partially obstructed airways in BOS lung tissue

Chymase, a serine protease released from mast cells, can be inhibited by SERPINA3 through the formation of a stable complex which impedes the enzymatic function of chymase^10^. Given this interaction, we were interested in the relationship between chymase and SERPINA3 expression by mast cells in the different airways in BOS lung tissue. We detected chymase-positive mast cells in connective tissue in normal airways (Figure 3A), whereas in partially obstructed airways we detected them in the connective tissue surrounding airway smooth muscle. In completely obstructed airways, these cells distributed to the connective tissue in the luminal obstruction. In fully obstructed airways we found a trend towards a higher area of chymase positivity compared to normal and partially obstructed airways (Figure 3B). To further investigate the interaction between chymase and SERPINA3, we investigated the extent of overlap between staining for chymase and SERPINA3. Surprisingly, we found only limited colocalization between the positive signals for chymase and SERPINA3 (Figure S4). We then questioned whether the binding between chymase and SERPINA3 was interfering with the detection by the anti-chymase antibody we applied. We therefore investigated protein-protein interactions, modelling the binding relationship between SERPINA3, chymase and the chymase antibody (Figure 3C). This prediction indeed indicated that SERPINA3 and chymase strongly interacted at a site that directly overlaps with the antibody binding site (figure 3C). This is indicated by the alignment of the blue reactive center loop in the SERPINA3 molecule and the chymase antibody binding region with the chymase molecule shown in green and orange. This modelling therefore suggested that our chymase antibody could not detect chymase when it was bound with SERPINA3. Consequently, the chymase that we did observe in airways was then not bound to SERPINA3 (and is referred to as unbound chymase from here on), indicating that we observed greater levels of unbound chymase in fully obstructed airways in BOS.

**Figure 3.**
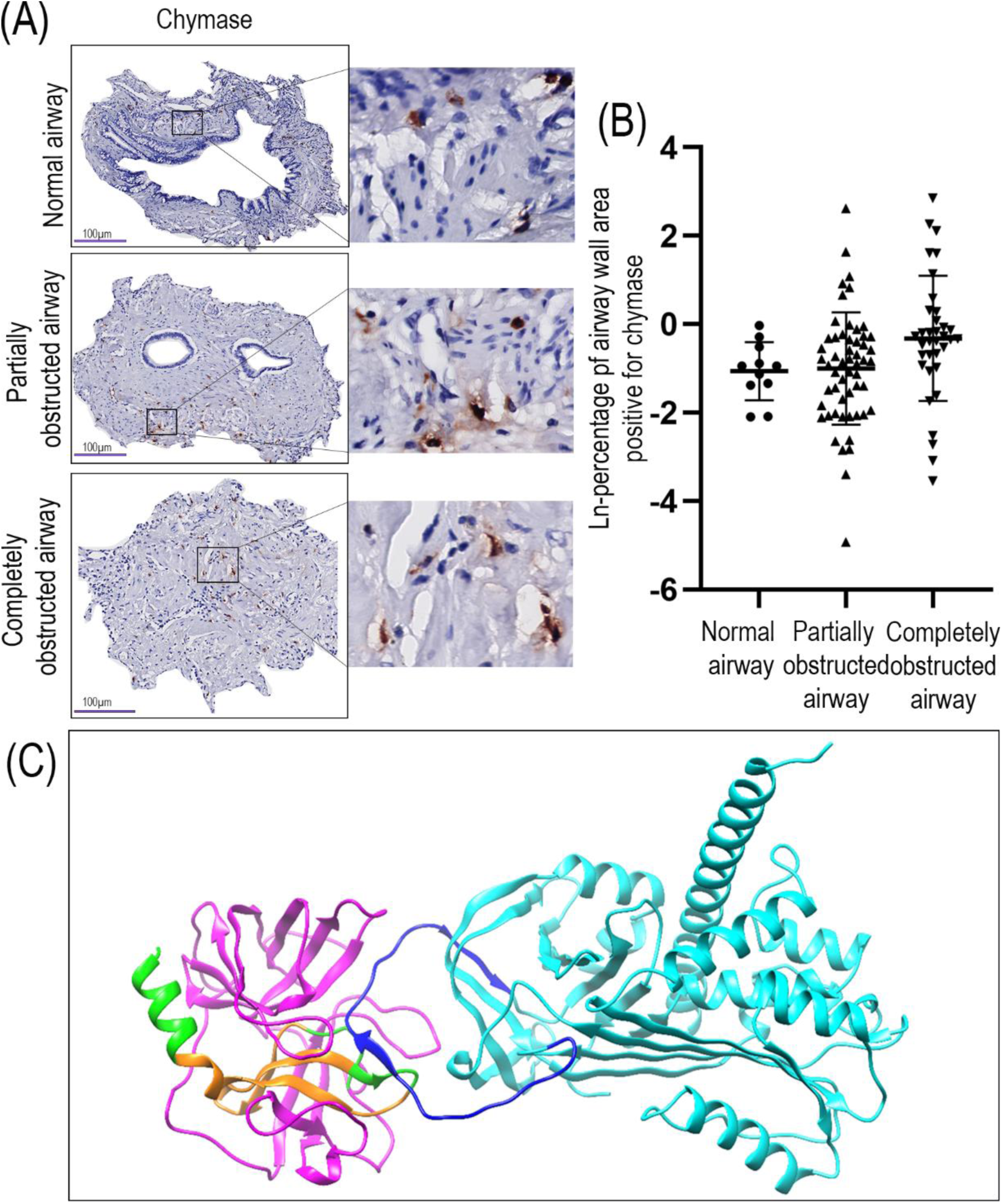
Localization of chymase-positive mast cells in airways of BOS lung tissue. Airways in lung tissue from patients with BOS stained for chymase using immunohistochemistry. Chymase expression is shown in red detected with NovaRed; nuclei are shown in blue detected with hematoxylin. Sections were imaged at the original objective magnification 400x and digital magnified images are shown next to each representative image. Three categories of airways were identified; normal, partially and completely obstructed airways. (A) Chymase-positive mast cells localized to normal airway, partially obstructed airway, and completely obstructed airway. (B) Percentages of chymase-positive area within normal, partially obstructed, and completely obstructed airways are presented as ln-transformed values and each datapoint represents an individual airway. Differences were analyzed using linear mixed effects regression. Data are presented as mean +/- SD, p<0.05 was considered statistically significant (n= 6 patients, number of airways per patient: 0-9 for normal airways, 0-21 for partially obstructed airways, 0-23 for completely obstructed airways). (C) AlphaFold model showing the interaction of SERPINA3 with chymase and the location of the chymase antibody binding site. SERPINA3 is shown in cyan with the reactive center loop of the molecule (amino acids (AA) 369-394) shown in blue. Chymase is represented in magenta, with the immunogenic peptide (AA 198-247) region shown in green, and the predicted epitopes for interaction with the antibody shown in orange (AA 204-219 and 224-237)

### Lower OPG deposition in partially and completely obstructed airways, which colocalized with SERPINA3 and tryptase in lung tissue from patients with BOS

Tryptase-positive mast cells can produce OPG protein^20^, a fibrosis associated factor. Moreover, chymase released from mast cells and OPG can negatively regulate each other under certain conditions^25^. We therefore wanted to investigate expression of OPG in airways of lung tissue from patients with BOS. Immunohistochemistry was again used to locate OPG in airways of those patients. OPG expression (Figure 4A) had a similar distribution pattern as seen for SERPINA3 and tryptase. We observed more OPG in normal airways compared to partially and completely obstructed airways (Figure 4B). Subsequently, we investigated if SERPINA3, tryptase and OPG colocalized in lung tissue from lung recipients with BOS. Using immunofluorescence, we found cells stained for SERPINA3, tryptase and OPG localized to the connective tissue of the airways.

**Figure 4.**
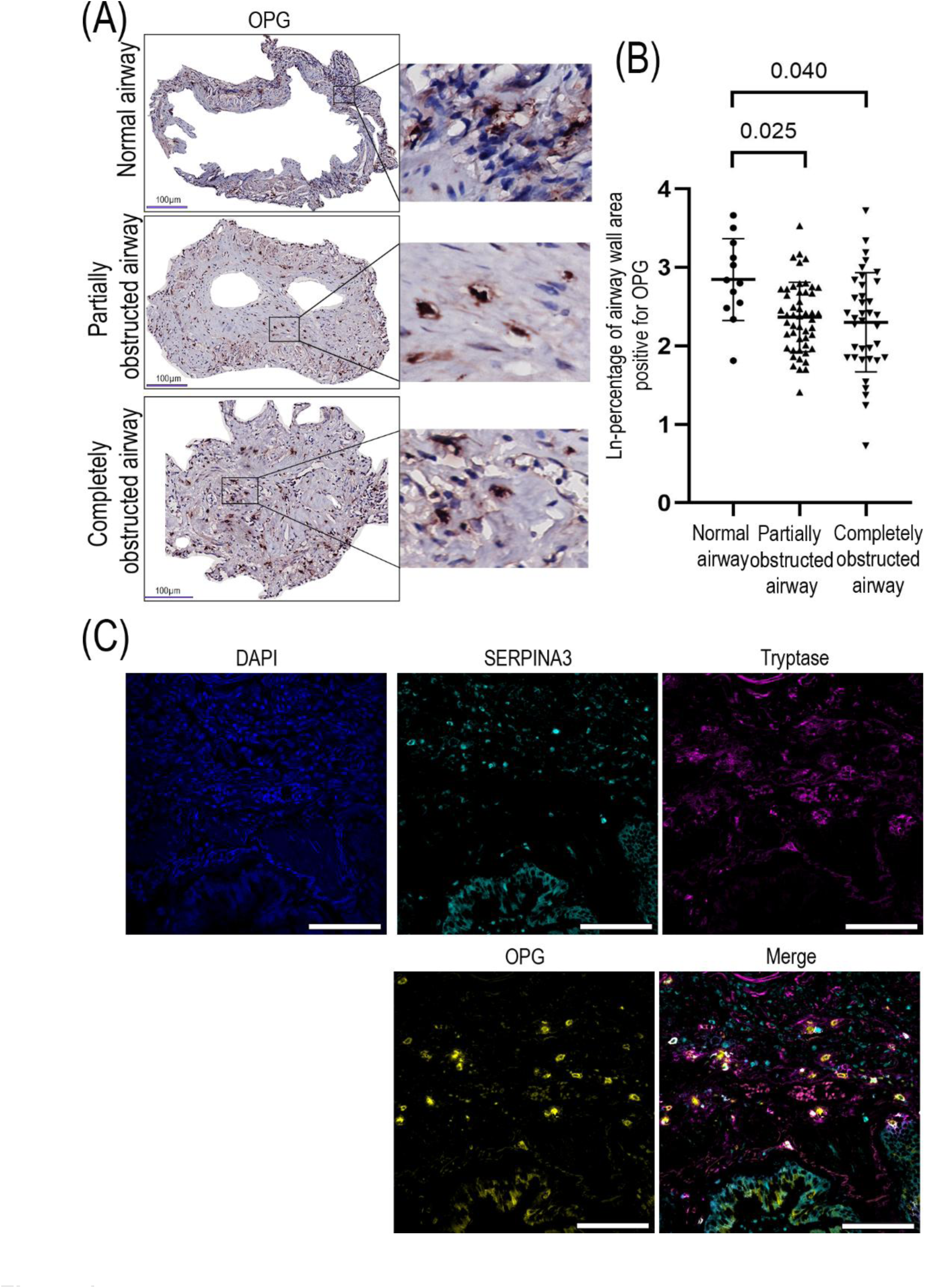
Localization of OPG in airway walls. Airways in lung tissue from patients with BOS were stained for OPG using immunohistochemistry. (A) OPG expression is shown in red detected with Nova Red; nuclei are shown in blue detected with hematoxylin. Sections were imaged at original objective magnification 400x, and digital magnified images are shown next to each representative image. (B) The percentage of OPG-positive area in normal, partially and completely obstructed airways is presented as ln-transformed values, and each datapoint represents an individual airway. Differences were analyzed using linear mixed effects regression. Data presented as mean +/- SD, p<0.05 was considered statistically significant (n= 6 patients, number of airways per patient: 0-10 for normal airways, 0-23 for partially obstructed airways, 0-26 for completely obstructed airways). (C) The colocalization of SERPINA3, tryptase and OPG in lung tissue from patients with BOS using immunofluorescence. Cell nuclei, SERPINA3, and tryptase are respectively shown in blue, cyan, and purple, colocalized expression is represented in white in the merged image (scale bar = 100µm).

### SERPINA3 and OPG may form a complex in the airways of lung tissue from patients with BOS

We were initially interested in whether SERPINA3, chymase, and OPG interacted in obstructed airways of patients with BOS. However, our modelling results showed that chymase could not be detected with our antibody when bound to SERPINA3. Consequently, we shifted our focus to exploring interactions between SERPINA3, tryptase (as an indicator of mast cells), and OPG.

A proximity ligation assay was used to investigate whether SERPINA3, tryptase, and OPG were located close enough within airway tissues to interact (Figure 5A). A strong proximity signal was detected between SERPINA3 and OPG and this signal associated with cells located within connective tissue of all airways. A more diffuse proximity signal was also observed in the extracellular matrix around cells in the connective tissue of partially obstructed airways and within luminal obstructions of completely obstructed airways. Similarly, a strong proximity signal between OPG and tryptase was detected, predominantly associated with cells within the connective tissue of both partially and completely obstructed airways. For SERPINA3 and tryptase, the proximity signal was weaker but displayed a distribution pattern similar to that of OPG and tryptase, suggesting that SERPINA3 and tryptase are potentially in close proximity.

**Figure 5:**
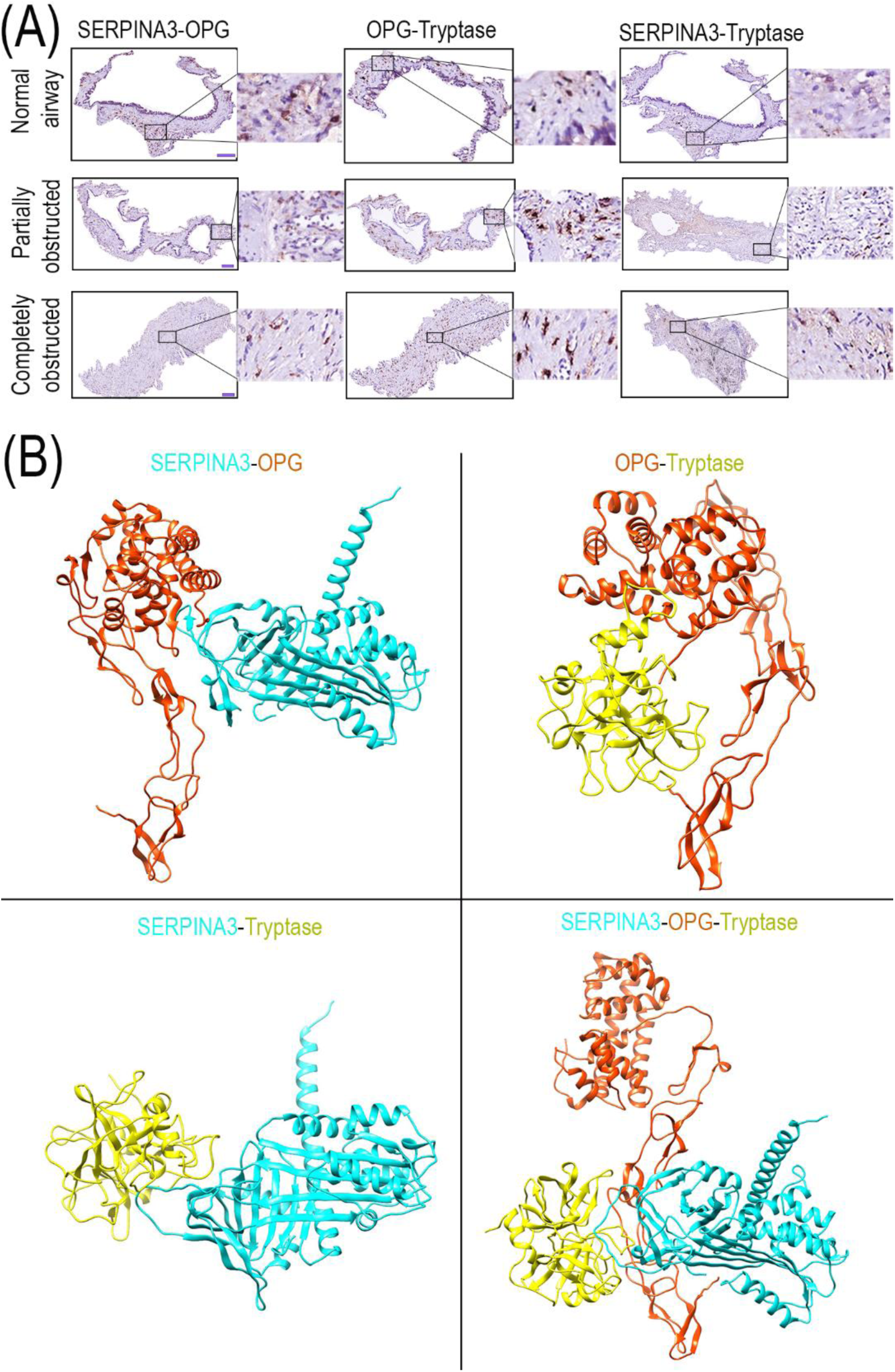
Proximity of SERPINA3, OPG, and tryptase in airways of lung tissue from recipients with BOS. (A) Representative images of proximity detection of SERPINA3 and OPG, OPG and tryptase, SERPINA3 and tryptase in lung tissue obtained from lung recipients with BOS. Each combination of two primary antibodies of SERPINA3, OPG and tryptase were simultaneously incubated with lung tissue. Positive signal (in red) is only found if distance between two proteins is within 40 nm^26^. Sections were imaged at original objective magnification 400x (bar = 100µm) and digital magnified images are shown next to each representative image. (B) AlphaFold models of interactions between SERPIN3A, OPG and tryptase. SERPINA3, OPG and tryptase are shown respectively in cyan, orange and yellow.

Next, we performed AlphaFold modelling again to predict interactions between SERPINA3 and OPG, OPG and tryptase, SERPINA3 and tryptase, as well as OPG, SERPINA3 and tryptase together (Figure 5B). This modelling showed a strong interaction between SERPINA3 and OPG, a moderate interaction between OPG and tryptase, a weak interaction between SERPINA3 and tryptase, and a potential interaction between SERPINA3, OPG and tryptase as inferred from the stabilizing energy of the different interactions (see table S8 in supporting information).

Altogether these data suggest that SERPINA3 and OPG may be released from mast cells and exhibit an interaction outside cells in the connective tissue of airways of BOS lung tissue.

### SERPINA3 and OPG are positively correlated in partially and completely obstructed airways in lung tissue from patients with BOS

Since a strong interaction between SERPINA3 and OPG was observed, we investigated whether their expression correlated in obliterated airways of patients with BOS. In normal airways, we found a correlation coefficient value between SERPINA3 and OPG of 0.77, which was not significant (supporting information Figure S5), which may have been caused by the small sample size of normal airways in our cohort. In partially and completely obstructed airways (Figure 6A-B), however, SERPINA3 and OPG expression exhibited a positive correlation. Recognizing SERPINA3 and OPG may be produced by tryptase-positive mast cells, we also examined the correlation between tryptase and SERPINA3, as well as tryptase and OPG. The data revealed no correlation between tryptase and SERPINA3, or tryptase and OPG, in normal or partially obstructed airways, (supporting information Figure S6). This is probably also due to the lack of power caused by the small sample size. In completely obstructed airways, however, we found a positive correlation between tryptase and SERPINA3, as well as between tryptase and OPG, in airways from patients with BOS. These data support our finding that OPG and SERPINA3 may be released from tryptase-positive cells and deposit in the connective tissue of airway walls in lung tissue from patients with BOS.

**Figure 6:**
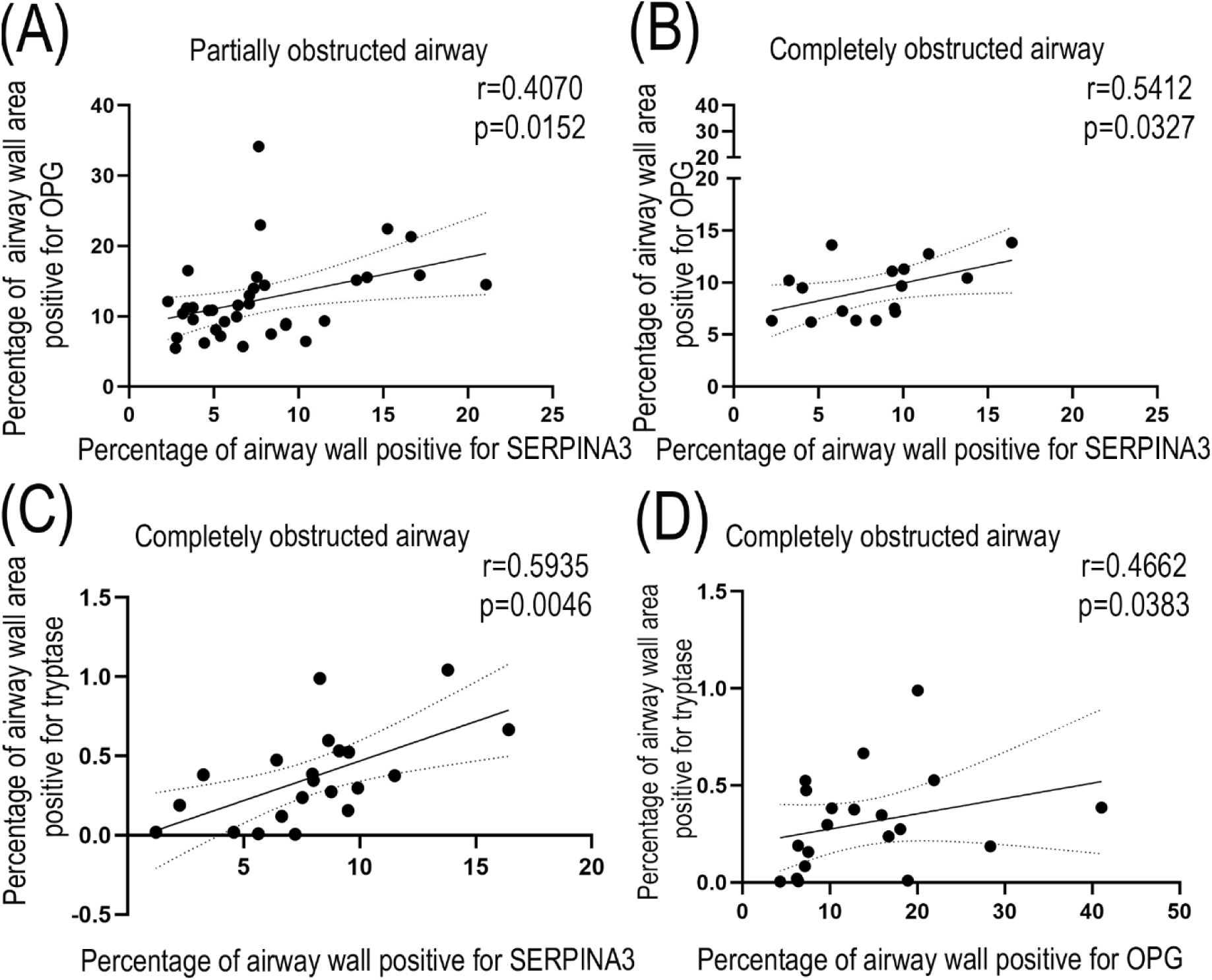
Correlations between expression of OPG and SERPINA3 or OPG and tryptase in partially and completely obstructed airways in lung tissue from recipients with BOS. (A) Correlation between SERPINA3 and OPG in partially obstructed airways (n= 35 airways). (B) Correlation between SERPINA3 and OPG in completely obstructed airways (n= 16 airways). (C) Correlation between SERPINA3 and tryptase in completely obstructed airways (n= 21 airways). (D) Correlation between OPG and tryptase in completely obstructed airways (n = 20 airways). Correlations were tested using a Spearman test, p<0.05 was considered significant.

## DISCUSSION

In this study, we investigated the presence of SERPINA3 and OPG and their potential interaction in airways of lung tissue from patients with BOS after lung transplantation. We found similar distribution patterns of SERPINA3 and OPG and we detected tryptase-positive mast cells in the submucosa of airways. Presence of these cells coincided with the cells positively stained for SERPINA3 and OPG, indicating SERPINA3 and OPG may be produced by tryptase-positive mast cells. Furthermore, we found that these two proteins were also close together in the extracellular space, suggesting tryptase-positive mast cells deposit them in the extracellular microenvironment. Overall, our findings suggest that SERPINA3 and OPG are likely released from tryptase-positive mast cells, and the association between the expression of SERPINA3 and OPG may be involved in airway obstruction in BOS.

The lower percentage SERPINA3 and OPG expression in partially and completely obstructed airways was likely a consequence of the increased amount of extracellular matrix present in the connective tissue and luminal obstruction in these abnormal airways. Similarly, the percentage area of tryptase-positive mast cells would also be impacted by the increase in extracellular matrix as obstruction of the airways increases. Therefore, the absolute amount of SERPINA3, OPG and tryptase may not have changed or may even be increased within these regions, however, the main localization of these proteins was different in BOS with less in the submucosa and more in the obliterating connective tissue in the region that was the lumen in obstructed airways. This differential localization suggests a role in the pathological process of BOS.

BOS has been recognized as a complex lung pathology with excessive extracellular matrix accumulation contributing to airway obstruction. Considering that our data suggest that SERPINA3 and OPG may be produced by tryptase-positive mast cells, which we found present around BOS airways, it is plausible to speculate that these two proteins are mediators playing a role in the process of mast cell-driven airway obstruction. During BOS, when the airways are partially obstructed, the resident tryptase-positive mast cells may migrate to the submucosa and connective tissue of the injured airways. Subsequently, SERPINA3 and OPG could be released from these cells. At advanced stages of airway obstruction, with high presence of SERPINA3 and OPG, these proteins may then diffuse to the connective tissue in the luminal obstruction. When SERPINA3 and OPG are released from tryptase-positive mast cells into the extracellular environment, these proteins may interact with fibrosis-associated proteins to regulate extracellular matrix remodeling. Clinically, inhibition of mast cells by for example montelukast (a leukotriene antagonist) started directly after lung transplantation could ameliorate the role of these mast cells, as well as SERPINA3 and OPG. A small prospective study performed with montelukast in progressive BOS patients^27^ did show attenuation of FEV1 decline but only for patients with BOS stage 1. The moment of starting montelukast could have influenced results, since irreversible fibrotic changes were already present in the later stages of BOS.

SERPINA3 has emerged as a novel fibrotic marker associated with various fibrotic disorders^28,29^. It plays a role as an essential protein in regulation of extracellular matrix gene expression in certain conditions. Matthew et al^14^ reported that silencing SERPINA3 resulted in low gene expression of collagen II, collagen IX and collagen XI in mesenchymal stem cells characterized by chondrogenesis. OPG is a glycoprotein shown to be involved in regulating extracellular matrix remodeling, for instance through its affinity for matrix proteins such as fibulin-1^30^ and connective tissue growth factor (CTGF)^31^. Both fibulin-1^32^ and CTGF^33^ have been recognized as important regulators of fibrotic processes. This knowledge implies that OPG may play roles in regulating extracellular matrix deposition, resulting in the airway obstruction in BOS. Given the proximity and the association, we observed between SERPINA3 and OPG in affected airways, we speculate that these two proteins may exhibit a synergistic relationship with each other in regulating obstruction of airways. An open question remains that needs further consideration: Do SERPINA3 and OPG released from tryptase-positive mast cells act as mediators to further stimulate the response of fibroblasts to produce more extracellular matrix in the injured airways of BOS?

An interesting puzzle in our study is the increase in both unbound chymase in lung tissue and SERPINA3 in serum of patients with BOS. The role of chymase is worth considering in the process leading to the development of airways obstruction, even though the chymase that was detected in this study was not in the form that was bound to SERPINA3, but rather the unbound chymase. SERPINA3 is an acute phase protein that is secreted by the liver in response to inflammation^12^. Therefore, the elevated level in serum could be coming from the liver. What the contribution of the lung is to circulating levels of SERPINA3 remains an open question. We noted that SERPINA3 was present in cells and the extracellular space in the submucosa and intraluminal of airways, where the percentage of unbound chymase detected per tissue area was greater in the completely obstructed airways, compared to unobstructed airways. A regulatory link between chymase in obstructed airway tissues and serum SERPINA3 in BOS remains elusive. We speculate that a feedforward signal provided by an increase in unbound chymase resulting from for example infection preceding BOS could enhance liver production of acute phase proteins including SERPINA3. In other cells chymase has been shown to negatively regulate OPG: when human osteoblasts were stimulated with chymase OPG protein was reduced ^25^. This potentially suggests that the high levels of unbound chymase in the obstructed airway may contribute to the downregulation of OPG expression in tryptase-positive mast cells. However, as the decreased proportion of OPG in completely obstructed airways was predominantly due to the increase in extracellular matrix driving an increase in tissue area, rather than a change in the apparent absolute amount of OPG detected in these airways, this seems unlikely. Further studies are needed to elucidate the impact of chymase on OPG in airway obstruction in BOS.

A limitation in this study is the unavailability of control explant tissue from lung transplantation patients without BOS. While all transplant patients undergo routine screening post transplantation, including surveillance biopsies, due to the patchy appearance of BOS and the size of surveillance biopsies from allografts, small airway analysis as described in this study can not be performed on such biopsies. In this study the unaffected airways in the explant tissue of BOS patients served as control, however for future studies, access to control explant tissue from LTx patients is preferred. Also, updated criteria by ISHLT (International Society for Heart & Lung Transplantation) for BOS staging have been implemented. However due to the retrospective nature of this study, guidelines available at the time of selection of serum samples were maintained for the duration of the study.

In conclusion, this study indicated SERPINA3 and OPG are likely produced by tryptase-positive mast cells, and they may form an extracellular complex in BOS airways with a possible role in the submucosal/intraluminal fibrosis. Further investigation is required to elucidate the precise mechanisms underlying obstruction of airways in BOS. An interesting focus could be how OPG and SERPINA3 influence extracellular matrix deposition. By identifying the interaction of OPG and SERPINA3, this study highlights a potential novel target for addressing the airway obstruction of fibrotic lesions in BOS lung disease.

