## Supplemental information for "A novel molecular interaction in bronchiolitis obliterans syndrome in lung transplantation patients: the role of SERPINA3 and osteoprotegerin"

**Appendix S1- METHODS**

**Patient information**

For serum analysis, lung transplant (LTx) patients who underwent bilateral lung transplantation between 2004 and 2017 in the University Medical Center Groningen were screened. BOS patients who progressed to stage three according to the International Society of Heart and Lung Transplantation Guidelines were selected if longitudinal serum samples were available. BOS patients were matched to non-BOS patients for sex, age at LTx, diagnosis necessitating LTx, immunosuppression and storage time of the samples (**Table S1**). Explants from LTx patients who received a re-transplantation due to BOS or deceased with end stage BOS were collected. All patients received immunosuppression according to protocol (**Table S2**). Patients provided written informed consent for use of material. The study was approved by the medical ethics committee of the University Medical Centre Groningen (METc 2014/077, METc 2021/610, research register number: 202000737), adheres to the UMCG Biobank Regulation and was conducted in accordance with the WMA Declaration of Helsinki and Declaration of Istanbul. Patients were enrolled in the ongoing, prospective TransplantLines Biobank and Cohort Study (ClinicalTrials.gov identifier: NCT03272841), in which, since June 2015, all (potential) solid organ transplantation patients and (potential) living organ donors (aged ≥18 years) at the University Medical Center Groningen (UMCG, The Netherlands) have been invited to participate.

**Table S1.** Patient characteristics of BOS and non-BOS patients for serum analysis

|  | BOS (n=19) | Non-BOS (n=19) | p-value |
| --- | --- | --- | --- |
| Patient sex, female (%) | 12 (63.2%) | 12(63.2%) | 1.0 |
| Age at LTx (y) | 55 [44-66] | 55 [43-66] | 0.6 |
| Underlying disease |  |  | 0.7 |
| COPD | 9† | 11† |  |
| CF | 2 | 2 |  |
| AATD | 5 | 3 |  |
| Scleroderma | 1 | 1 |  |
| Other | 2* | 2* |  |
| Donor sex, female (%) | 12(63.2%) | 11(57.9%) | 1.0 |
| Donor age | 52± 10 | 46 ± 12 | 0.1 |
| Donor pack years | 2.5 [0-12.5] | 0 [0-15] | 0.8 |
| Immunosuppression after LTx |  |  | 1.0 |
| Tacrolimus/azathioprin/prednisolon | 5 (26.3%) | 5(26.3%) |  |
| Tacrolimus/mmf/prednisolon | 14 (73.7%) | 14(73.7%) |  |
| Patients with documented diagnosis of  viral infection after LTx | 10 (52.6%) | 3 (15.8%) | 0.08 |
| Time to diagnosis of BOS (y) | 2.8 [1.9-5.8] | NA |  |
| Pulmonary function test - in L (n)** |  |  |  |
| FEV1 BOS stage 1 | 1.97±0.64 (18) | 2.78±0.39 (18) | <0.001 |
| FVC BOS stage 1 | 3.18±0.81 (18) | 3.81±0.89 (18) | 0.1 |
| FEV1 BOS stage 2 | 1.59±0.53 (18) | 2.76±0.47 (8) | <0.001 |
| FVC BOS stage 2 | 3.03±0.88 (18) | 3.57±0.93 (8) | 0.17 |
| FEV1 BOS stage 3 | 1.21±0.39 (19) | 2.70±0.46 (16) | <0.001 |
| FVC BOS stage 3 | 2.58±0.81 (19) | 3.74±0.89 (16) | <0.001 |
| FEV1% predicted BOS stage 1 | 71.50±15.06 (18) | 95.50±24.14 (18) | <0.001 |
| FVC% predicted BOS stage 1 | 90.5±16.30 (18) | 106.00±18.59 (18) | 0.006 |
| FEV1% predicted BOS stage 2 | 53.33±13.08 (18) | 105.25±20.21 (8) | <0.001 |
| FVC% predicted BOS stage 2 | 84.28±15.49 (18) | 110.88±15.35 (8) | <0.001 |
| FEV1% predicted BOS stage 3  FVC% predicted BOS stage 3 | 40.0±8.98 (19)  70.00±14.56 (19) | 98.00±19.34 (16)  107.00± 12.55 (16) | <0.001  <0.001 |

† 2 BOS patients with alpha-1 antitrypsin deficiency were matched to non-BOS COPD controls. *One BOS patient with histiocytosis X matched with a non-BOS patient with pulmonary fibrosis. One BOS patient with pulmonary fibrosis after infection matched with non-BOS patient with selective IgG2 deficiency and bronchiectasis. **Non-BOS patients (n=19) were matched to BOS patients (n=19) for time after transplantation regarding pulmonary function tests performed. Abbreviations: BOS: bronchiolitis obliterans syndrome; COPD: chronic obstructive pulmonary disease; CF: cystic fibrosis; AATD: alpha-1 anti-trypsin deficiency; LTx: lung transplantation, y: years; m: months; n: number; MMF; mycophenolic acid, FEV1: forced expiratory volume in one second; FVC: forced vital capacity; FEV1% predicted: forced expiratory volume as a percentage of the predicted value; FVC% predicted: forced vital capacity as a percentage of the predicted value.

**Table S2.** Patient characteristics of BOS patients (n=6) for tissue analysis

|  | BOS (n=6) |
| --- | --- |
| Patient sex, female (%) | 1 (16.7%) |
| Age at first LTx (y)  Time until re-LTX or death (y) | 35 [20 - 48]  6.1 [4.0-10.7]† |
| Underlying disease |  |
| CF | 2 (33.3%) |
| AATD | 4 (66.7%) |
| Donor age | 45 [29-50] |
| Donor pack years | 0 [0-3] |
| Pulmonary function test - L* |  |
| FEV1 BOS stage 1 | 2.4 [1.9 - 2.8] |
| FVC BOS stage 1 | 4.1 [3.6 - 4.8] |
| FEV1 BOS stage 3 | 1.4 [1.1 - 1.9] |
| FVC BOS stage 3 | 3.1 [2.9 - 3.7] |
| FEV1% predicted BOS stage 1 | 68.0 [51.8-72.3] |
| FVC% predicted BOS stage 1 | 82.5[78.0-90.0] |
| FEV1% predicted BOS stage 3 | 42.5 [33.0 - 46.0] |
| FVC% predicted BOS stage 3 | 69.5 [67.3 - 88.3] |

† 5 BOS patients underwent re-transplantation because of BOS, one patient was included after obduction. *Pulmonary function testing of 4 patients included. Missing values due to first transplantation and follow up in other centers. Abbreviations: Abbreviations: BOS: bronchiolitis obliterans syndrome; CF: cystic fibrosis; AATD: alpha-1 anti-trypsin deficiency; LTx: lung transplantation, y: years; n: number; L=liters; FEV1: forced expiratory volume in one second; FVC: forced vital capacity.

**Hematoxylin & Eosin (HE) staining**

Three μm sections of paraffin-embedded lung tissues from patients with BOS were deparaffinized in xylol for 2 times 10 min. Subsequently, the sections were washed two times with 100% ethanol, with 96% ethanol, with 70% and with demi water each for 10 seconds. Then hematoxylin was stained in the slides for 10 min. After washing the slides with tap water and incubating the slide in tap water for 10 min, the slides were stained with eosin for 2 min. Next, the sections were dehydrated with 96% ethanol, and with 100% each for 10 seconds. Slides were air-dried for 10 min before mounting the slides with mounting medium (SP15100, Thermo Fisher Scientific, United States). Finally, the slides were scanned at 40x magnification using slide scanner Hamamatsu Nanozoomer (Hamamatsu Photonic K.K., Japan).

**Verhoeff staining**

Three μm sections of paraffin-embedded lung tissues from patients with BOS were deparaffinized in xylol for twice for 10 min. The sections were then incubated in demi water to remove residual xylene. Subsequently, slides were incubated in Verhoeff working solution for 15 min, followed by a brief rinse in demi water. Next, slides were incubated with 1% ferric chloride for 1 minute, followed by wash slides with demi water for three times. Then slides were incubated in van Gieson’s stain solution for 2 min and wash them with demi water after staining. Dehydrated slides in 100% alcohol, air-dried out and mounting slides with mounting media (Thermo Fisher Scientific, United States). Slides were scanned at 40x magnification using slide scanner Hamamatsu Nanozoomer (Hamamatsu Photonic K.K., Japan), reagents used in Verhoef staining are summarized in Table S3.

**Table S3.** Reagents used in Verhoeff staining

| Reagent | Composition |
| --- | --- |
| Verhoeff Stock Solution (A) | 5 g hematoxylin in 100 mL 100% ethanol |
| Verhoeff Stock Solution (B) | 5 g Ferric Chloride in 50ml demi water |
| Verhoeff Stock Solution (C) | 1 g potassium iodide in 2.5 mL demi water, then add 0.5 g iodine. |
| Verhoeff Working Solution | 25 mL Solution A + 10 mL Solution B + 10 mL Solution C |
| Acid fuchsin | 0.1 g acid fuchsin in 10 mL distilled water |
| Van Gieson’s solution | 6 mL saturated picric acid + 1 mL 1% acid fuchsin |
| Ferric Chloride | 2 g in 100 ml demi water |

**ELISA**

Serum SERPINA3 level was measured by Human alpha 1-Antichymotrypsin ELISA Kit (ab157706, Abcam, Cambridge, United Kingdom), sample dilution is 1: 5000 optimized according to the manufacturer's instructions.

**Immunohistochemical staining**

Three μm sections of paraffin-embedded lung tissues from patients with BOS were deparaffinized and antigens were retrieved through incubating in 10 mM citrate buffer at pH 6.0 at 100℃ for 15 min. Endogenous peroxidases were blocked in a solution of PBS containing 0.3% hydrogen peroxide (H2O2) (Merck KGaA, Darmstadt, Germany) at room temperature for 30 min. After washing three times in 1X Phosphate Buffered Saline (PBS), lung tissue sections were blocked with 4% bovine serum albumin (BSA) in PBS at room temperature for 30 min. Next the sections were incubated with primary antibodies overnight at 4℃, then incubated with secondary antibodies for 45 min at room temperature (**Table S3**), before being washed with PBS three times and subsequently with demi water three times. Finally, the staining was visualized by incubating the slides for 10 minutes with Nova Red staining solution (SK-4800, BRUNSCHWIG CHEMIE, Amsterdam, Netherlands). All sections were counterstained using hematoxylin, before being dehydrated and mounted with histological mounting media (Thermo Fisher Scientific, United States). Slides were scanned at 40x magnification using slide scanner Hamamatsu Nanozoomer (Hamamatsu Photonic K.K., Japan).

**Table S4.** Antibodies using in immunohistochemical staining

| Primary  antibody | Primary  antibody  source | Primary  antibody dilution | Secondary  antibody | Secondary antibody  source | Secondary antibody dilution |
| --- | --- | --- | --- | --- | --- |
| Anti-AACT | ab180492, Abcam | 1:1000 | Goat anti-rabbit immunoglobulin-HRP | Dako P0488, Dako, Amsterdam, NL | 1:100 |
| Anti-mast cell Tryptase | ab2378, Abcam | 1:100000 | Rabbit anti-mouse immunoglobulin-HRP | Dako P0260, | 1:100 |
| Anti-chymase | JB74-32,  Novus Biologicals, Abingdon,  United Kingdom | 1:16000 | Goat anti-rabbit immunoglobulin-HRP | Dako P0488 | 1:100 |
| Anti-osteoprotegerin | ab183910, Abcam | 1:300 | Goat anti-rabbit immunoglobulin-HRP | Dako P0488 | 1:100 |

Abbreviations: AACT: alpha-1-antichymotrypsin

**Immunofluorescent staining**

For simultaneous detection of SERPINA3, OPG and Tryptase sections of formalin-fixed paraffin embedded lung tissue were sequentially stained for each protein with tyramide signal amplification. This method allows the detection of multiple primary antibodies without species cross-reactivity and is based on an antibody stripping protocol which removes primary antibodies and secondary antibodies but not the visualization signal. Sections of formalin-fixed paraffin embedded lung tissue were deparaffinized, incubated with citrate buffer (10mM sodium citrate, pH 6) for 15 min at 100^o^C for antigen retrieval and cooled down for 30 min at room temperature. The slides were washed with Tris Buffered Saline (TBS) and endogenous peroxidase was blocked by incubating the slides in TBS containing 0.3% hydrogen peroxide for 30 min at room temperature. The slides were incubated with rabbit anti-SERPINA3 (Abcam, ab180492, 1:500) in TBS with 1% BSA for 1 hour at room temperature followed by goat anti-rabbit horseradish peroxidase-conjugated secondary antibody (DAKO, P0448, 1:100). Visualization was performed using tyramide signal amplification process with Opal 570 Reagent Pack (AKOYA Biosciences, FP1488001KT), slides were incubated in Opal 570 Reagent Pack diluted 1:200 in 0.1M Borate buffer containing 0.003% H_2_O_2_ for 10 min at room temperature. Next slides were incubated again in citrate buffer (10mM sodium citrate, pH6) for 15 min at 100^o^C to remove all antibodies and cooled down for 30 minutes at room temperature. The slides were then incubated with mouse anti-Tryptase (Abcam, ab2378, 1:10000) and rabbit anti-OPG (Abcam, ab183910, 1:300) in TBS with 1% BSA for 90 min at room temperature followed by goat anti-rabbit horseradish peroxidase-conjugated secondary antibody (DAKO, P0448, 1:100) and donkey anti-mouse Alexa647-conjugated secondary antibody (Thermo, A-31571, 1:500). Visualization for OPG was performed using tyramide signal amplification process with Opal 520 Reagent Pack (AKOYA Biosciences, FP1487001KT), slides were incubated in Opal 520 Reagent Pack diluted 1:200 in 0.1M Borate buffer containing 0.003% H_2_O_2_ for 10 min at room temperature. Negative staining controls were performed by omitting the primary antibody incubations and by staining for only 1 protein but including all secondary steps for the other proteins. Nuclei were visualized with DAPI and slides were mounted in citifluor (Thermo, 17970-025). Fluorescent images were acquired using a Zeiss LSM780 confocal scanning microscope (Carl Zeiss, Oberkochen, Germany) and an Olympus VS200 fluorescent slide scanner (Olympus Corporation, Tokyo, Japan).

**Proximity ligation assay**

The proximity ligation assay was performed with the Nave Bright-HRP kit (NB.MR.HRP.100, Navinci, Uppsala, Sweden). The kit enables chromogenic visualization of protein-protein interactions. Sections of formalin-fixed paraffin embedded lung tissue were deparaffinized and subsequently incubated with citrate buffer (10mM sodium citrate, pH 6) for 15 min at 100^o^C for antigen retrieval. The slides were cooled down for 30 min at room temperature and washed with TBS. Endogenous peroxidase was blocked by incubating the slides in TBS containing 0.3% hydrogen peroxide for 30 min at room temperature. The slides were washed with TBS and then incubated overnight with 4 different combinations of primary antibodies: mouse anti-SERPINA3 (Proteintech, 66078-1—Ig, 1:100) + rabbit anti-OPG (Abcam, ab183910, 1:300), mouse anti-tryptase (Abcam, ab2378, 1:10000) + rabbit anti-OPG (Abcam, ab183910, 1:300) and mouse anti-tryptase (Abcam, ab2378, 1:10000) + rabbit anti-SERPINA3 (Abcam, ab180492, 1:1000). Also, incubations with only 1 primary antibody were taken along to assess background staining. All antibodies were diluted in TBS containing 2% BSA. After washing with TBS 0.1% TWEEN® 20 (P1379, Sigma-Aldrich, Amsterdam, Netherlands), the slides were incubated with anti-mouse Navenibody and anti-rabbit Navenibody in Navenibody diluent at 37 °C for 1 hour, followed by enzymatic ligation, rolling PCR amplification, HRP incubation and substrate development according to the manufacturer’s instructions (Navinci, NB.MR.HRP.100). Nuclei were visualized with Mayer’s hematoxylin (Sigma Aldrich). Slides were scanned using a Hamamatsu NanoZoomer 2.0 HT digital scanner (Hamamatsu Photonics).

**Airway categorization and identification**

Slides from recipients with BOS were scanned and uploaded into Qupath^1^, version 0.5.0. Airways were identified in the H & E, Verhoef and MSB stained slides. All airways were identified by an experienced lung pathologist (WT) (summarized in **Figure S1**). All airways without presence of cartilage and submucosal glands, were included in the image analyses in this study. Airways that were excluded were the last part of terminal bronchiole going into respiratory bronchiole (so where the airway smooth muscle cell layer is interrupted), because of the absence of ECM**.** All airways were identified and categorized according to severity of obliteration (shown in **Table S5**). In this study, we focused on three airway categories including normal airway, partially obstructed airway and completely obstructed airway.

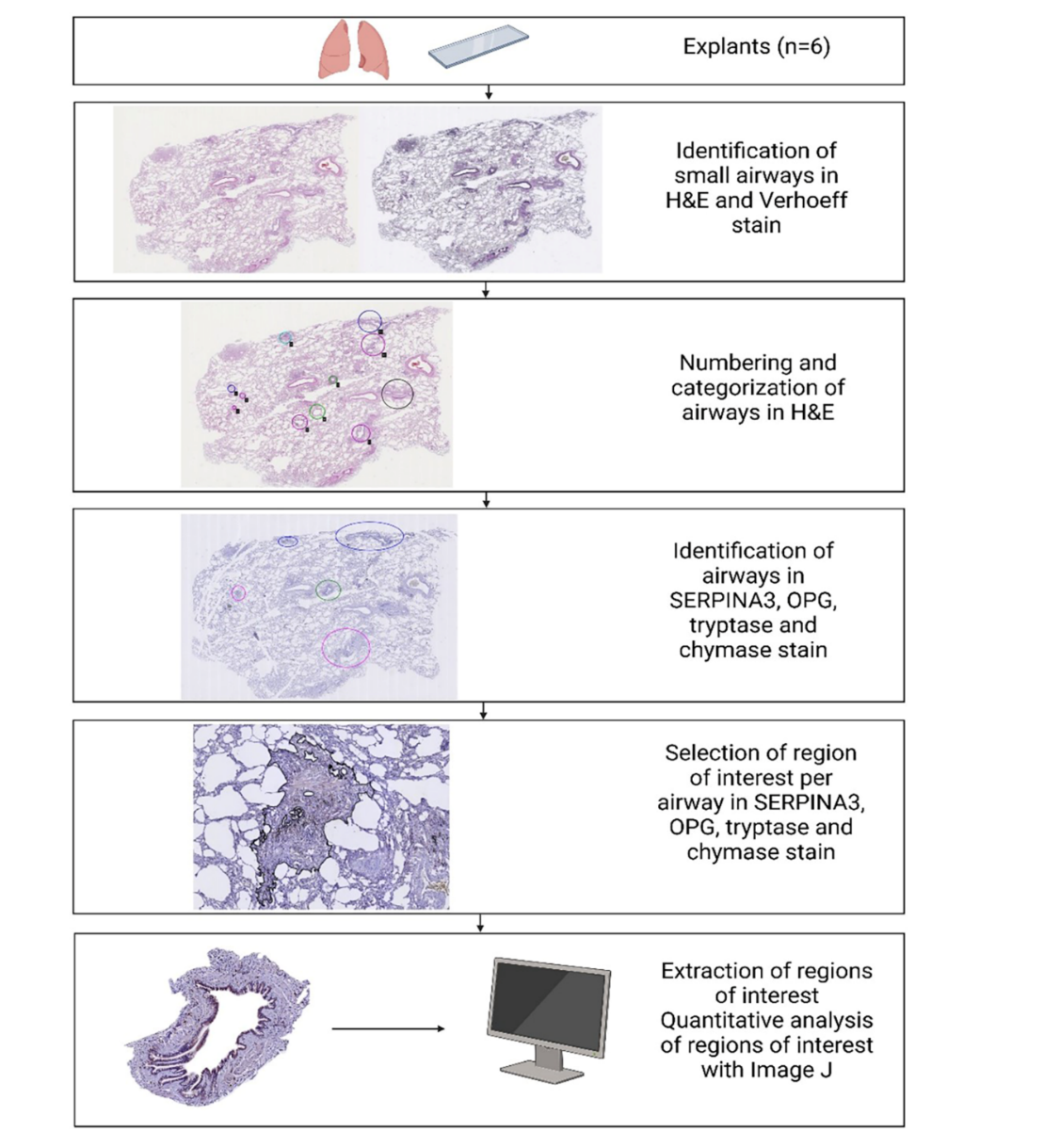

**Figure S1**. Flow diagram of identification of airway, airway categorization and selection of region of interest. Explant tissue (n=6) was stained with hematoxylin and eosin (H&E), as well as Verhoef stain. Airways were identified in H&E and Verhoeff stain. Airways were numbered and categorized into different airway categories (normal, partially obstructed, completely obstructed). Airways were also identified in SERPINA3, tryptase, chymase and OPG stained sections and categorized according to the classification code established in the H&E-stained section. Region of interest defined as the outer border of the adventitia of the airway to the inner border of the airway lumen for completely, partially and normal airways, was identified and selected. Regions of interest were extracted and quantitative analysis was performed using Image J. Abbreviations: n: number; H&E: hematoxylin and eosin; SERPINA3; SERPINA family member A3; OPG: osteoprotegerin.

**Table S5.** Airway categorization and identification

| Categorization | Identification |
| --- | --- |
| Normal airway | Unaffected airways. |
| Partially obstructed, non-active airway | Airway with fibrotic obliteration with a greater distance between the airway smooth muscle layer and the epithelial layer than normal airway, with at least a partially recognizable lumen. |
| Completely obstructed airway | Airways that have a completely obstructed lumen. |

**Airways of interest annotation**

The region of interest was defined as the outer border of the adventitia of the airway to the inner border of the airway lumen for partially and normal airways. When airways were completely obstructed, the circumference of the outer border of the adventitia was identified as the boundary of the region of interest. Aberrations within the images that were excluded from the area of interest are described in **Figure S2** and **Table S6**.

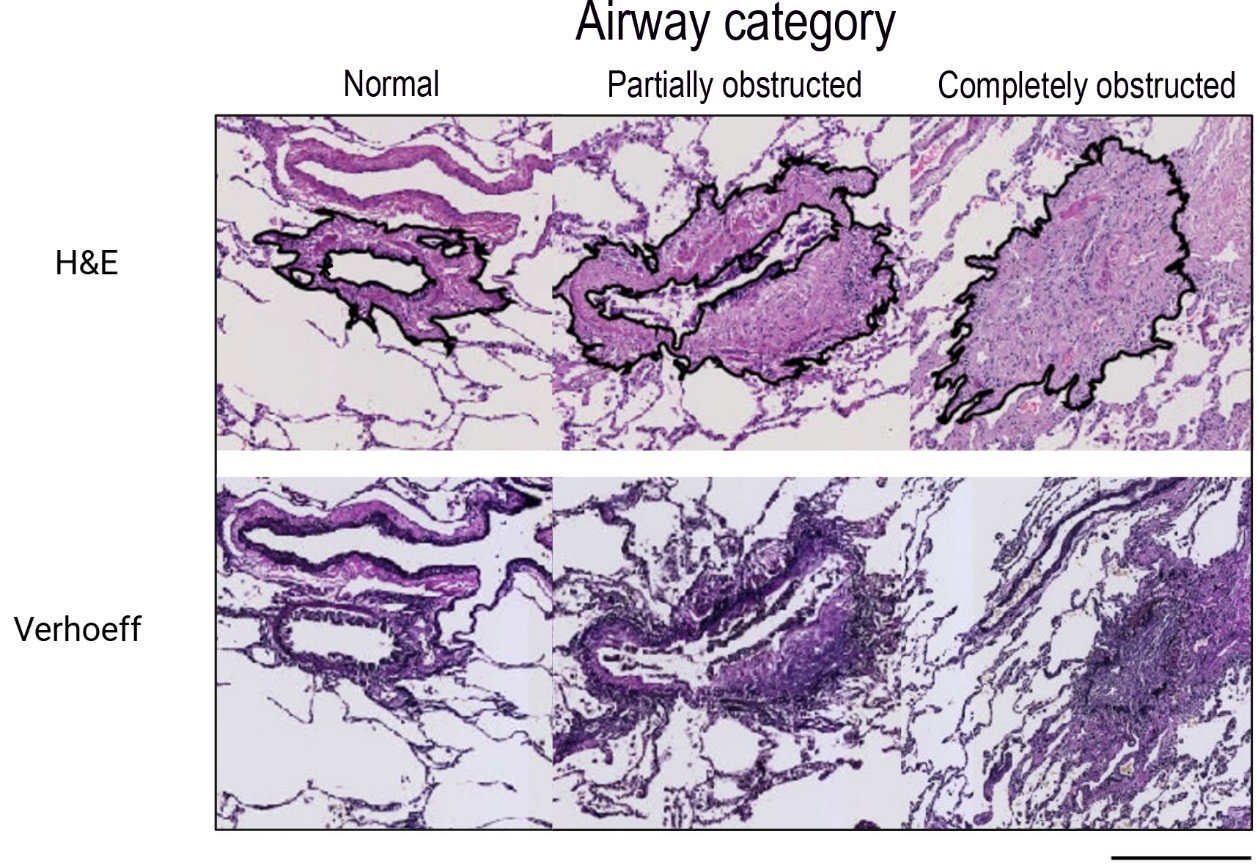

**Figure S2**. Airway categorization and identification of region of interest per airway category. Upper row shows airway categories in hematoxylin and eosin, bottom row shows airway categories in Verhoeff stain. Black line indicates region of interest, defined as the outer border of the adventitia of the airway to the inner border of the airway lumen for partially and normal airways. When airways were completely obstructed, the circumference of the outer border of the adventitia was identified as the boundary of the region of interest. Definition of airway categories: normal; airway not affected by obliterative bronchiolitis, partially obstructed: airway with fibrotic obliteration with a greater distance between the airway smooth muscle layer and the epithelial layer, with at least a partial recognizable lumen; completely obstructed: airways that have a completely obstructed lumen. Magnification 40x, black scale bar 400 µm. Abbreviations: HE; hematoxylin and eosin stain.

**Table S6.** Features excluded from the regions of interest

| **To be excluded area** | **Details** |
| --- | --- |
| Lumen area in unaffected and partially obstructed airways | No cells present; not contributing to airway surface containing extracellular matrix. |
| Arteriole in Adventitia with Diameter >50 µm | Exclude if no clear arteriole wall; if wall is identifiable, exclude both lumen and wall. |
| Larger Vessels | Border determined by matrix change or muscle layer of vessel. |
| Epithelial Layer Loose from Basal Membrane | If completely loose, exclude both epithelium and lumen; if still attached, include. |
| Protrusions of Airway with Inflammatory Infiltrate | Excluded due to the little extracellular matrix present in these areas and small protrusions without immune |

**Image analyses**

A summary of the sample size analyzed for each staining is shown in **Table S7**. For analyses of staining images, briefly, the NDPI files were firstly converted to TIF images in Aperio ImageScope (v12.3.3.5048, Leica) and subsequently processed in Adobe Photoshop 2024 (Adobe Inc. California, United States) to exclude artificial areas such as folded tissues and carbon-based pigments. Next, Image J win 64 software was utilized to quantify the area with positive staining. All images were split into blue (hematoxylin-image), and red (NovaRed-image) pixels using color deconvolution plugin of Image J^2^. To calculate the total amount of tissue, images were converted to 8-bit grey scale. Total number of pixels representing total tissue area versus positively stained tissue area were identified using the threshold feature of Image J. Percentage of positive area within the whole region was calculated using the formula below. Data analyses were performed with R software 4.4.0 (Boston, Massachusetts, USA).

$$Area \left( \% \right)=\frac{Number of pixels positive for NovaRed}{total Number of pixels positive for tissue}*100$$

**Table S7.** Summary of sample size analyzed for each staining

| Staining | Patient (n) | Block/  Patient (n) | Normal airway (n) | Partially obstructed airway (n) | Completely obstructed airway (n) |
| --- | --- | --- | --- | --- | --- |
| SERPINA3 | 6 | 2-5 | 9 | 46 | 33 |
| Tryptase | 6 | 2-5 | 8 | 56 | 29 |
| Chymase | 6 | 2-5 | 11 | 55 | 35 |
| OPG | 6 | 2-5 | 13 | 51 | 38 |

**Prediction of biomolecular interactions**

AlphaFold Server^3^(<https://alphafoldserver.com>) was used to predict biomolecular interactions. The strength of the predicted protein-protein interactions was assessed using PPCheck^4^ (<https://caps.ncbs.res.in/ppcheck/>). Open Babel^5^ version 3.1.1 hosted on cheminfo.org was used to convert structure file formats. Molecular graphics images were produced using UCSF Chimera^6^ version 1.17. Prediction of antigenic peptides was performed using the Universidad Complutense Madrid, Immunomedicine Group tool (<http://imed.med.ucm.es/Tools/antigenic.pl>) using the method of Kolaskar and Tongaonkar^7^.

**Table S8.** Analysis of Biomolecular interactions

| Proteins | Pdb/Uniprot | ipTM | pTM | H Bond  kJ/mol | Electro  static kJ/mol | Van der Walls kJ/mol | Total stabalizing  kJ/mol |
| --- | --- | --- | --- | --- | --- | --- | --- |
| Chymase  SERPINA3 | 1KLT  3DLW | 0.8 | 0.63 | -122.85 | 15.81 | -369.94 | -476.98 |
| SERPINA3  Chymase | P01011 (24-423) P23946 (22-247) | 0.79 | 0.64 | -110.99 | -7.42 | -316.79 | -435.2 |
| SERPINA3  Chymase | P01011 (1-423) P23946 (1-247) | 0.76 | 0.64 | -104.31 | -17.49 | -321.57 | -443.37 |
| SERPINA3  OPG | P01011 (24-423) O00300 (22-401) | 0.17 | 0.5 | -71.79 | -72.36 | -220 | -364.15 |
| SERPINA3  OPG | P01011 (1-423) O00300 (1-401) | 0.16 | 0.48 | -56.95 | -19.57 | 149.87 | 73.35 |
| OPG  Tryptase | O00300 (22-401) Q15661 (31-275) | 0.18 | 0.47 | -28.88 | .17 | -247.63 | -276.35 |
| OPG  Tryptase | O00300 (1-401) Q15661 (1-275) | 0.51 | 0.55 | -27.81 | -73.14 | -170.03 | -270.98 |
| SERPINA3  Tryptase | P01011 (24-423) Q15661 (31-275) | 0.42 | 0.61 | -92.2 | -9.06 | 2.31 | -98.95 |
| SERPINA3  Tryptase | P01011 (1-423) Q15661 (1-275) | 0.42 | 0.57 | -112.47 | -91.28 | -287.59 | -491.34 |
| Tryptase (a)  SERPINA3 (b)  OPG (c) | Q15661 (31-275) P01011 (24-423)  O00300 (22-401) | 0.3 | 0.46 | -88.91  -7.44    -18.91 | -115.82    -11.52    -38.78 | -322.8  -66.55  -31.49 | -527.53 a/b  -85.51 a/c  -89.18 b/c |
| SERPINA3 (a)  OPG (b)  Tryptase (c) | P01011 (1-423) O00300 (1-401)  Q15661 (1-275)  O00300 (1-401) | 0.27 | 0.44 | -184.76  -124.59  0 | -93.94  -91.73  -5.86 | 2263.94  -365.10  -12.02 | 1985.24 a/b  -581.42 a/c  -17.87 b/c |

**Figure S3.** Longitudinal mean ± standard deviation of LN transformed serum SERPINA3 in non-BOS patients (n=19) and in BOS patients (n=19) over different timepoints.

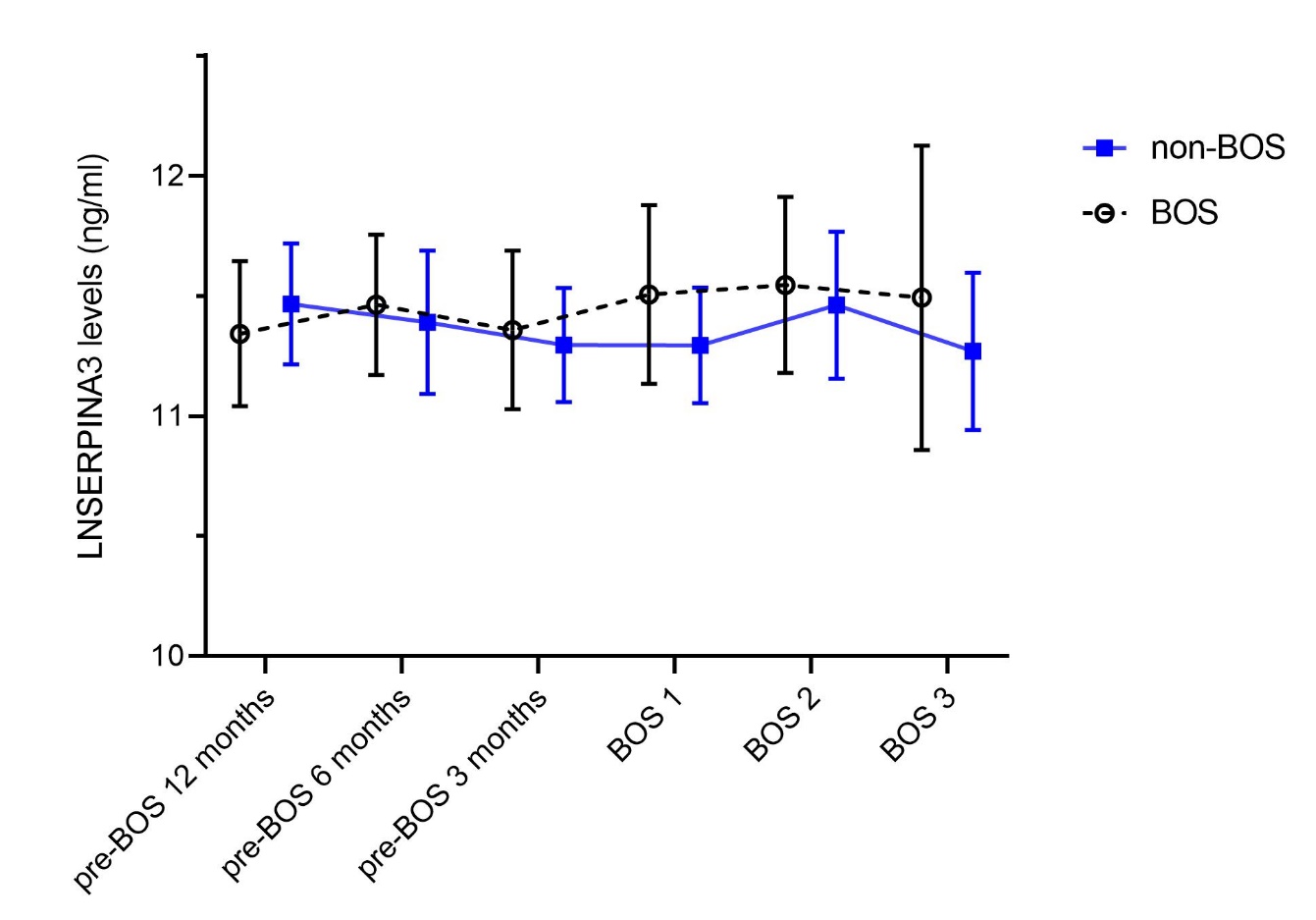

**Figure S3**. Longitudinal mean ± standard deviation of LN transformed serum SERPINA3 levels in ng/ml in non-BOS patients and BOS patients over different timepoints. The solid line represents the non-BOS group, the dotted line represents the BOS group. Error bars indicate the standard deviation. Abbreviations: BOS; bronchiolitis obliterans syndrome.

**Figure S4.** Colocalization of SERPINA3 and chymase in lung tissue from patients with BOS.

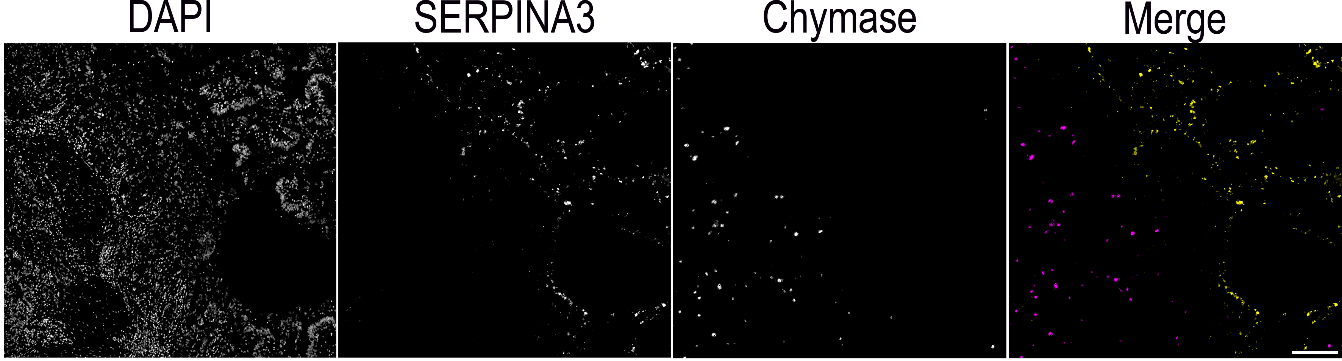

**Figure S4:** showed the colocalization of SERPINA3 and chymase in lung tissue from patients with BOS. FFPE lung tissue sections from patients with BOS stained with SERPINA3 and chymase using immunofluorescence. SERPINA3, and chymase are respectively shown in yellow, and purple (scare bar = 100µm).

**Figure S5.** Correlation between SERPINA3 and OPG in normal airway in lung tissue from patients with BOS

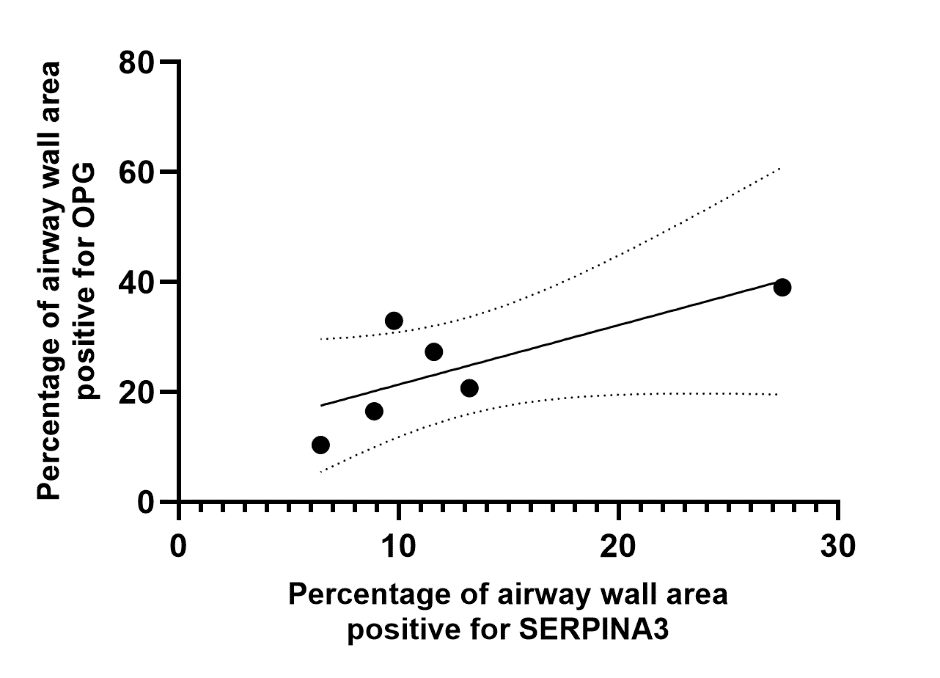

**Figure S5:** Correlation between SERPINA3 and OPG in normal airways (n= 6 airways, r=0.7714, p= 0.1028). Correlations were calculated using a Spearman test.

**Figure S6.** Correlation between tryptase and, SERPINA3, tryptase and OPG in normal and partially obstructed airways in lung tissue from patients with BOS

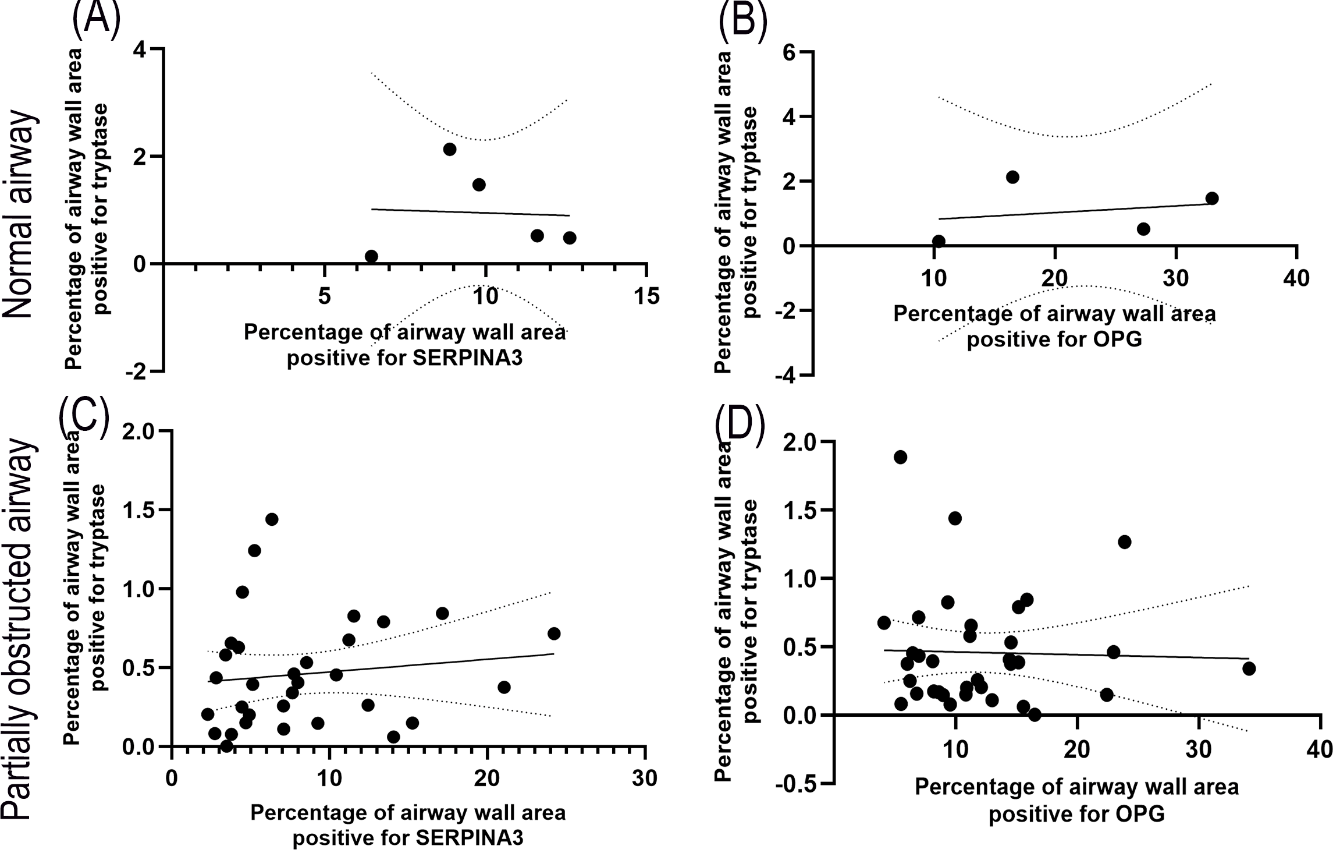

**Figure S6:** Correlation between tryptase and SERPINA3, tryptase and OPG in normal airways and partially obstructed airways. (A) Correlation between tryptase and SERPINA3 in normal airway (n=5, r= 0.000, p>0.999), (B) Correlation between tryptase and OPG in normal airway (n=4, r= 0.400, p=0.7500), (C) Correlation between tryptase and SERPINA3 in partially obstructed airway (n= 32, r= 0.2093, p= 0.2503), (D) Correlation between tryptase and SERPINA3 in partially obstructed airway (n= 35, r= -0.04062, p= 0.8168). Correlations were calculated using a Spearman test, r=0.7714, p= 0.1028.
